# Comparative regulomics provides novel insight into the evolution of wood formation across dicot and conifer trees

**DOI:** 10.1101/2025.06.28.661522

**Authors:** Eduardo Rodriguez, Siri Birkeland, Ellen Dimmen Chapple, Samuel Fredriksson, Zulema Carracedo Lorenzo, Teitur Ahlgren Kalman, Vikash Kumar, Jason Hill, Hannele Tuominen, Ewa J. Mellerowicz, Nathaniel R. Street, Torgeir R. Hvidsten

**Author notes:** These authors contributed equally to this work (Eduardo Rodriguez, Siri Birkeland, Ellen Dimmen Chapple).

## Abstract

Understanding the regulatory program underlying wood formation is key to improving biomass production and carbon sequestration in trees. However, how wood formation evolved and how these programs have been rewired across lineages remains unclear. Here, we present the first high-spatial-resolution evo-devo resource spanning the wood transcriptomes of six tree species - three dicots and three conifers - capturing 250 million years of evolutionary divergence. Using orthology-aware co-expression network analysis, we identified genes with conserved and lineage-specific expression patterns. By integrating chromatin accessibility data and transcription factor motif analysis, we further inferred regulatory networks for xylem differentiation and secondary cell wall formation. We demonstrate how this dataset can be used to answer long standing questions in wood biology related to differences in acetylation of cell wall polymers and master regulators of xylem specification across dicot and conifer tree species. The data offer a foundational resource for the tree biology and evo-devo communities, and are publicly available at PlantGenIE.org.

## Introduction

Trees are an essential component of terrestrial ecosystems and wood is one of the planet’s largest sources of renewable biomass^1,2^. Beyond their ecological and economic importance, trees also play a key role in carbon sequestration, with wood serving as a long-term reservoir for storing the carbon fixed through photosynthesis^3^. In fact, forests comprise the second-largest carbon store on Earth after the oceans^4^. However, the regulatory mechanisms underlying wood formation remain poorly understood - especially the extent to which these mechanisms are shared or derived across diverse tree lineages. Bridging this knowledge gap is essential for uncovering fundamental principles of vascular development and for guiding efforts to enhance biomass production and long-term carbon capture.

Wood, or secondary xylem, is not only a fundamental structural and functional component of trees but also a model for studying the molecular mechanisms determining cell differentiation and tissue development. Wood formation arises from the activity of the vascular cambium, a narrow layer of secondary meristematic cells that produce secondary phloem towards the outer layers of the stem (i.e. towards the protective bark) and secondary xylem inwards^5^. Cambial cells differentiate by cell enlargement followed by secondary cell wall (SCW) deposition. Although *Arabidopsis thaliana* (Arabidopsis) is an herbaceous plant, it does exhibit limited wood formation^6^ and has therefore been studied to provide a foundation for understanding the gene regulatory networks underlying wood formation in trees^7^. In Arabidopsis, these networks consists of several NAC (NAM, ATAF1/2 and CUC2) domain transcription factors (TFs) that control specific MYB (myeloblastosis) family TFs, which in turn regulate genes essential for SCW formation such as SCW cellulose synthase (CesA) genes^8^.

While the fundamental wood cell types and processes are shared by most trees, significant differences exist between the two major tree-containing plant lineages: gymnosperms, of which conifers are the most diverse and well-studied group, and angiosperms^9^. Conifer wood, often referred to as softwood, is primarily composed of tracheids, which transport water and provide mechanical support. In contrast, angiosperm wood, or hardwood, evolved two additional types of xylem elements that serve specific functions; the vessel elements being dedicated to water transport and the fibers providing structural support^10^. The chemical composition of cell wall polymers also differs between hard and soft wood types. For instance, lignin is generally less abundant in hardwood fibers than in softwood cells and differs in monomer composition: angiosperm lignin contains both guaiacyl (G) and syringyl (S) units (derived from coniferyl and sinapyl alcohols, respectively) with minor amounts of p-hydroxyphenyl (H) units, while conifer lignin contains predominantly G units with low amounts of H units, and typically lacks S units^9^.

Comparative genomics studies have suggested that regulatory changes play a key role in determining tree growth habit and wood formation^7^. To uncover the molecular mechanisms underlying the wood phenotype, several studies have applied high-spatial-resolution ‘omics technologies, including transcriptomics^11–14^, proteomics^15^, metabolomics^13,16^ and epigenomics^17^. More recently, single-cell transcriptomics has provided additional insight into the cellular complexity and developmental dynamics of wood-forming tissues^18,19^.

To fully uncover how wood formation evolved and how developmental programs have been rewired across lineages, evolutionary developmental biology (evo-devo) approaches are essential. Evo-devo studies have the potential to identify how changes in gene regulation contributed to the emergence and diversification of wood formation^20^. While evo-devo has previously been used to, for example, compare gene expression between normal, tension and opposite wood^21,22^, this approach has not yet been applied using high-spatial-resolution data to perform comparative regulomics. In addition, the regulatory mechanisms underlying differences between angiosperm hardwood and gymnosperm softwood have never been compared. A comparative analysis of angiosperm and conifer species offers a powerful approach to identify key regulatory and developmental differences, as well as conserved components of the gene regulatory network underlying wood formation in trees. In this study, we leverage data on accessible chromatin and transcription factor binding with high-spatial-resolution transcriptomics data across wood-forming tissues in three dicots (the group of angiosperms that form wood via a bifacial vascular cambium) and three conifers to deepen our understanding of gene regulation during secondary growth, and to shed light on the evolutionary and functional dynamics of vascular tissue development in trees.

## Results and Discussion

We present a resource of high-spatial-resolution transcriptomes across dicot and conifer wood formation. In the following sections, we (1) reveal conserved and diverged gene expression patterns between the two lineages, (2) use these findings to answer selected long-standing questions in wood biology and (3) compare dicot and conifer gene regulatory networks.

### High-spatial-resolution wood transcriptomics across dicot and conifer tree species

We first generated and analyzed high-spatial-resolution RNA sequencing (RNA-Seq) data spanning the secondary phloem, vascular cambium, and wood-forming tissues of three dicots (aspen/*Populus tremula*^11^, birch/*Betula pendula* and cherry/*Prunus avium*) and three conifers (Norway spruce/*Picea abies*, Scots pine/*Pinus sylvestris* and Lodgepole pine/*Pinus contorta*). For each species, three to four replicate trees were sampled from clonal copies of a single genotype, with all replicates being mature trees growing in the field. First, 15-µm thick longitudinal tangential cryosections were obtained across developing phloem and wood-forming tissues. These cryosections were then pooled into 25-28 samples per replicate tree: single sections were used within the cambial meristem region, pools of three sections representing expanding and secondary cell wall (SCW)-forming xylem, and pools of nine sections capturing maturing xylem (Tab. S1)^11^. Finally, gene expression datasets were created by performing RNA-seq on the pooled samples (522 samples in total).

By mapping RNA-seq reads to five reference genomes (both pine species were mapped to the Scots pine genome sequence as no reference is available for lodgepole pine), and estimating relative gene expression values for each gene, we obtained six transcriptomes with 11,000-20,000 genes expressed across 489 samples (Tab. S2). Gene expression patterns during different stages of wood formation were highly structured and consistent across all six species, as shown by unsupervised principal component analysis (Fig. 1A) and unsupervised hierarchical clustering (Fig. S1) - both of which largely reconstructed the sampling order based on gene expression similarity. Hierarchical clustering further revealed that all six transcriptomes underwent major reprogramming events that largely coincided with (a) the dividing cambial cells located between phloem and xylem differentiation, (b) the end of cell expansion and the beginning of SCW formation, and (c) the end of SCW deposition (Fig. S1). This was supported by anatomical observations (Tab. S1) and the distinct expression profiles of five well-characterized marker genes (Fig 1B)^11^.

**Fig. 1.**
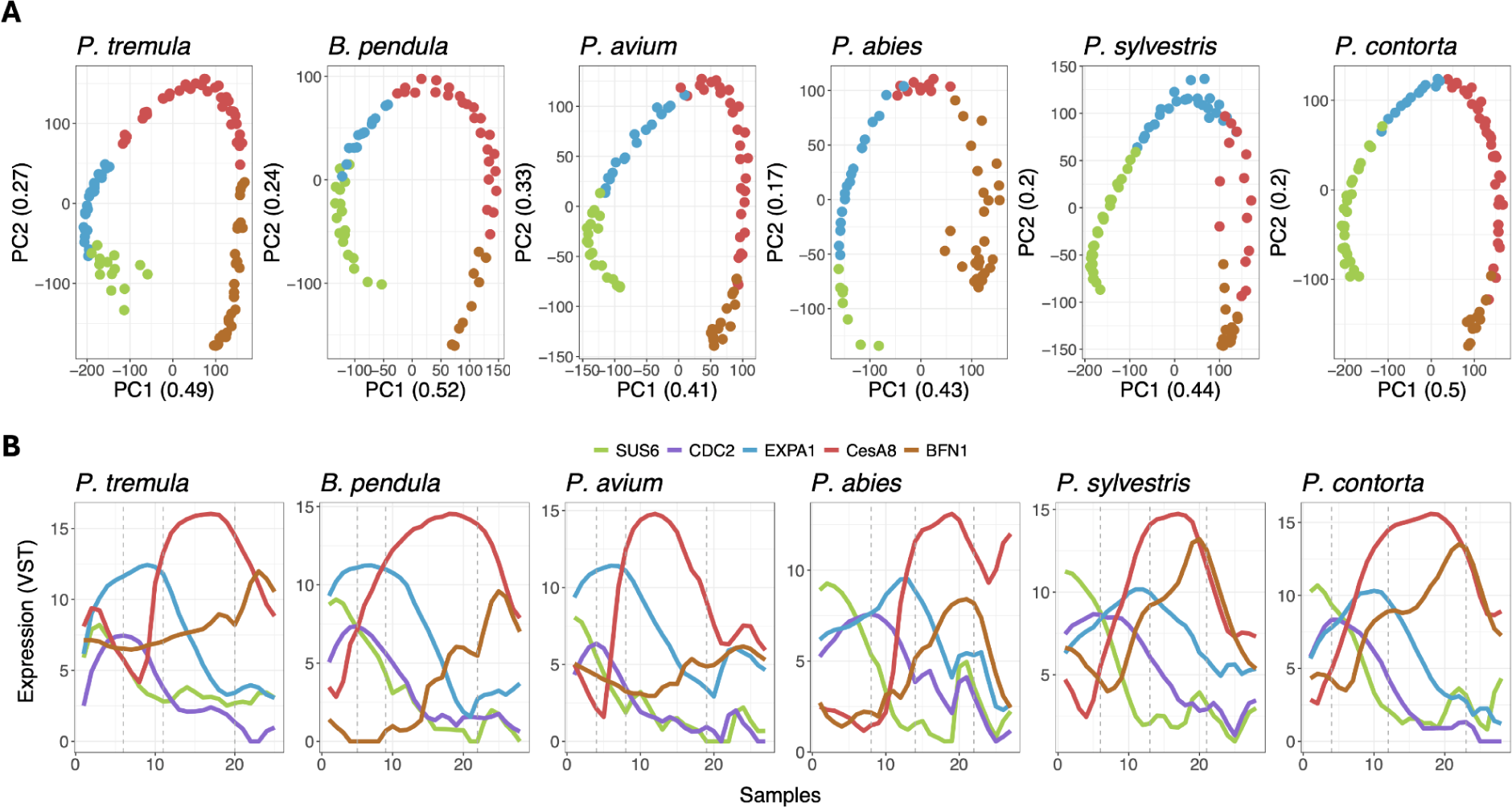
**Principal component analysis (PCA) and genes marking the different stages of wood formation**. (A) PCA of all samples across the replicate trees colored by stages of wood formation. Percent explained variance in parentheses. (B) Expression profiles of genes marking the phloem (SUS6), cambial activity (CDC2), the expanding xylem (EXPA1), the SCW-forming xylem (CesA8), and mature xylem/cell death (BFN1). Dotted lines mark, from left to right, the middle of the cambium, the end of the expansion zone and the end of the SCW zone.

### Comparative wood transcriptomics

To identify genes with conserved and diverged expression, we first identified ortholog groups (orthogroups) across 27 plant species based on protein sequence. This resulted in 5,564 orthogroups with genes expressed in wood in all six species, as well as 2,093/3,649 uniquely expressed orthogroups in dicots/conifer wood, respectively (Fig. 2A). Based on ortholog identification, we then compared gene expression across wood forming tissues. The sample series were not directly comparable among species due to anatomical and growth differences. We therefore inferred co-expression networks for each species and identified co-expressologs; pairs of orthologous genes from two different species that are embedded in conserved co-expression relationships. Specifically, two genes are considered co-expressologs if they are orthologs and are co-expressed with other genes that are also orthologs of one another across species^23^ (Fig. 2B). This definition emphasizes conserved network context rather than direct expression similarity, in contrast to the original definition of an expressolog^24^. Since co-expression typically only occurs between genes with smooth, developmentally regulated expression profiles, and these profiles need to be shared by a significant number of genes for a co-expressolog to be identified, this approach will not identify constitutively expressed genes or genes with rare expression profiles (typically genes with random spikes of expression) as co-expressologs. We identified approximately 3,000-10,000 orthogroups with at least one co-expressolog per species pair (Tab. S3), with a clear phylogenetic signal of higher numbers of within- than between-lineage co-expressologs (Fig. S2A).

**Fig. 2.**
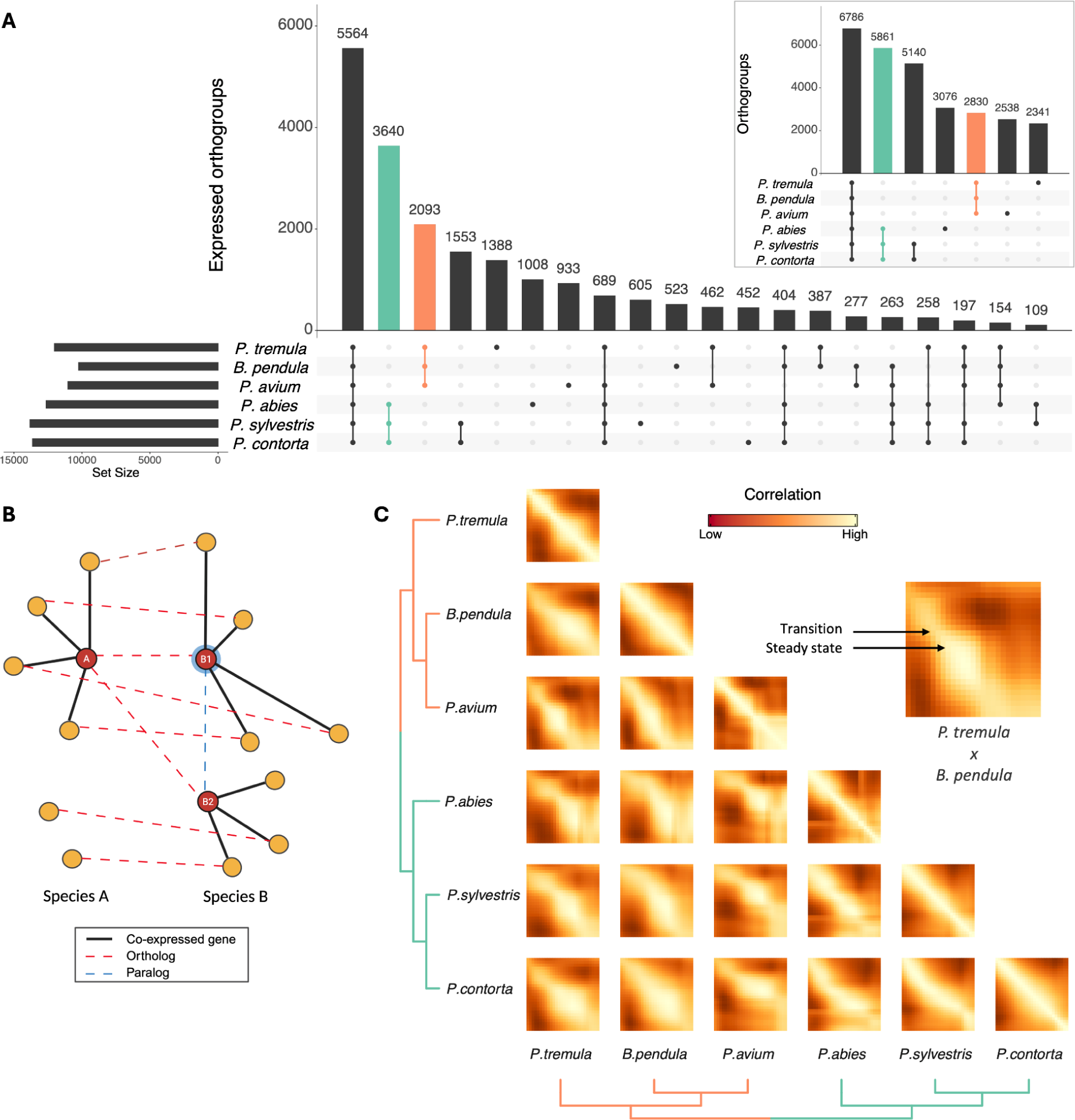
Comparisons of six wood transcriptomes. (A) Expressed orthogroups: Upset plot showing the number of orthogroups with genes expressed in wood for each species (horizontal bars) and the number of orthogroups with expressed genes from different subsets of species (vertical bars). Orthogroups: Upset plot with all genes (right corner). (B) Method for identifying orthologs with conserved co-expression (co-expressolog). For each ortholog pair, a p-value is computed that reflects to what degree their co-expressed genes also are orthologs. In this example, A and B1 are co-expressologs while A and B2 are not. (C) Heatmaps displaying the correlation of samples across species pairs, computed using co-expressologs. Only one replicate tree is shown per species. Colors are scaled for each heatmap (each species pair).

Using co-expressologs, we next analyzed the similarity among the six wood transcriptomes. This revealed that gene expression during wood formation is largely conserved across 250 million years of evolution (Fig. 2C): samples from the same developmental stage were more similar to each other across species than samples from different stages within species (diagonals in Fig. 2C and bimodal distribution of correlations in Fig. S2B). This analysis also showed a clear phylogenetic signal, with corresponding samples within dicots and conifers showing much higher correlation (∼0.75) than samples compared between the two lineages (∼0.55; Fig. S2B). Our analyses also revealed the same major stages of wood formation, and the corresponding reprogramming events, seen in cluster analysis of genes (Fig. S1) (i.e. yellow squares of steady state expression in Fig. 2C, of which the SCW stage was the most apparent, separated by transitions/reprogramming events). This observation is consistent with a suggested modular nature of secondary growth^25^.

### Genes with conserved wood co-expression across 250 Myr of evolution

To identify gene sets with conserved expression across all six species, we analyzed co-expressolog networks constructed within each orthogroup. In these networks, nodes represent genes and edges connect orthologous gene pairs that are co-expressologs, reflecting conserved co-expression relationships across species. We then searched for fully connected subnetworks, or cliques, containing exactly one gene from each species (Fig. 3A). This resulted in 70,458 cliques from 2,098 orthogroups, of which 2,145 cliques were non-overlapping (unique) (Tab. S4). Hence, 38% of the expressed orthogroups (2,098/5,564, Fig. 2A) could be classified as having consistently conserved expression across all species. Since co-expressologs are based on conserved co-expression and not expression profile similarity (Fig. 2B), these cliques could, in principle, reveal shifts in expression profiles, for example between dicots and conifers. However, the set of 2,145 unique cliques displayed strikingly similar expression profiles across the six species, where genes typically showed expression specific to the same stages of wood formation across all species (Fig. 3B). Conserved cliques covered major stages and processes of wood formation, as evident from expression profiles (Fig. 3B) and gene function enrichment analysis (Tab. S5).

**Fig. 3.**
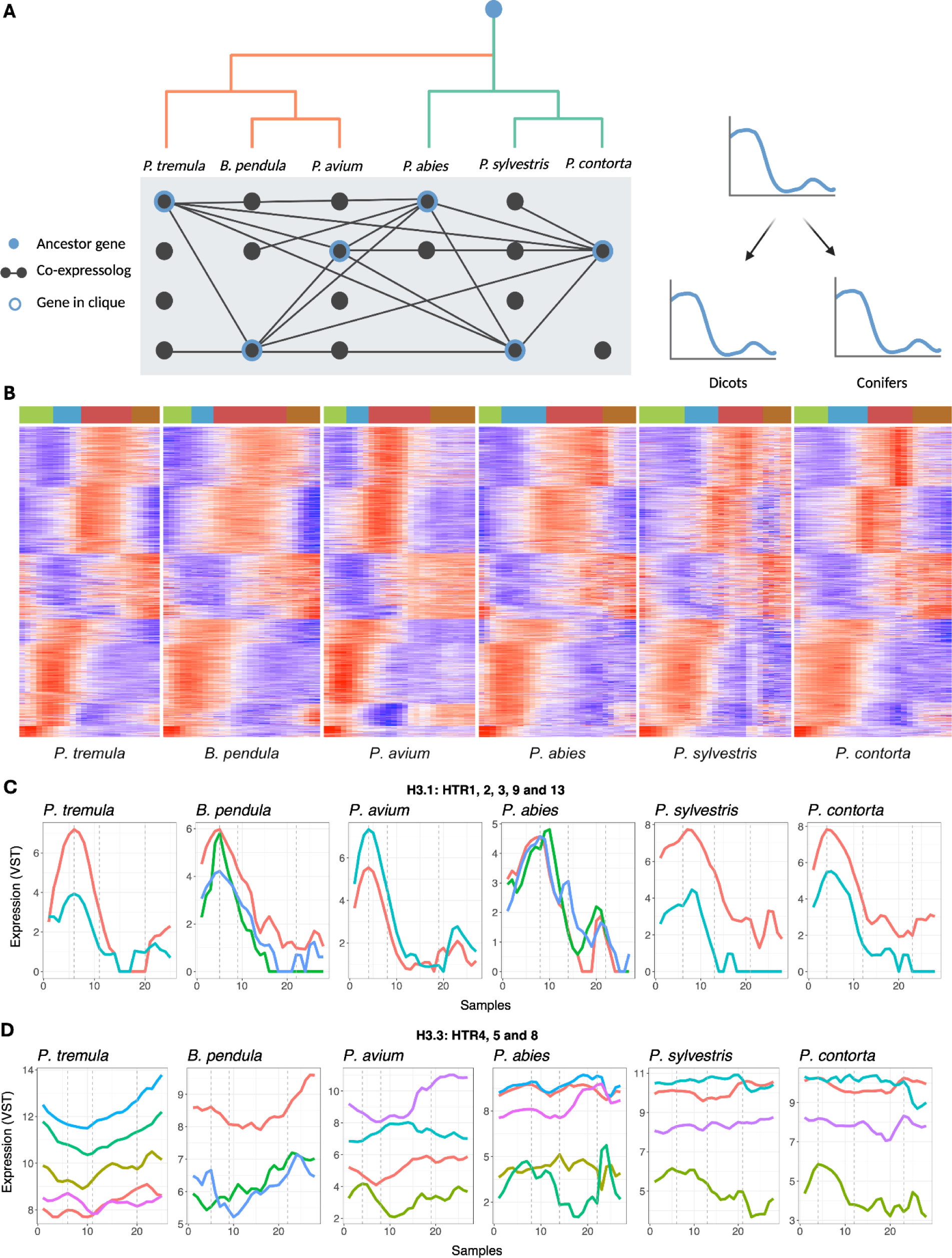
Genes with conserved co-expression across all species. (A) Conceptual figure of how we selected cliques of genes with conserved co-expression across all six species. A network was created for each orthogroup, connecting genes that are co-expressologs. Fully connected subnetworks with one gene from each species (i.e. cliques) were identified. Note that cliques from the same orthogroup can partly overlap. (B) Heatmap of unique cliques (i.e. cliques with no genes in common). Each row is a clique of six genes, one from each species. Each column is a sample from one replicate tree from each species, and the top color bar indicates the wood formation stages as established by hierarchical clustering. (C,D) One orthogroup (OG0000095) contained all Arabidopsis H3.1 (HTR1, 2, 3, 9 and 13) and H3.3 (HTR4, 5 and 8) genes. The gene tree is available in Fig. S3. (C) Genes in top-cliques from the H3.1-clade of the orthogroup gene tree, showing conserved expression across all species. (D) All expressed genes in the H3.3-clade of the orthogroup gene tree. H3.1 is known to either lack introns (H3.1) or contain introns (H3.3) across the plant tree of life^28^, and genes not meeting these criteria were therefore removed.

The species we studied have long divergence times, and the orthogroups typically contain multiple genes from each species and have gene phylogenies that are difficult to interpret, with genes often segregating into distinct dicot-specific and conifer-specific clades. Moreover, they often contain a large number of conifer genes due to extensive local duplications in these species^26^. These challenges can be addressed using cliques, which identify orthologs with similar co-expression patterns across clades, helping to pinpoint the most likely ‘functional’ orthologs. To illustrate this, we studied the orthogroup of H3.1 and H3.3 histones. H3.1 is incorporated during DNA replication and is thus linked to proliferating cells, while H3.3 is incorporated into chromatin in response to transcriptional activity, maintaining chromatin accessibility^27^. The gene tree of the histone orthogroup separated H3.1 and H3.3 histones into different clades, but it was not possible to establish 1-1 orthology between the characterized Arabidopsis genes and genes in the other species within these clades (Fig. S3). However, we found a number of H3.1-cliques containing genes with clear expression peaks in the cambium of all six species (Fig. 3C). Hence, H3.1 marks the proliferative phase of wood formation, where cambial cells are undergoing mitotic divisions before differentiating into xylem or phloem, indicating that DNA replication is largely restricted to the cambium during wood formation. On the other hand, we did not identify any H3.3-cliques as these genes showed constitutive expression across the wood series, with no developmentally regulated expression profiles (Fig. 3D). This suggests ongoing chromatin remodeling in developing phloem and xylem, and indicates a role of epigenetic regulation in major transcriptome reprogramming events.

### Genes with diverged expression between dicots and conifers during wood formation

Dicot and conifer wood have very different vascular structures and SCW chemical composition. To identify genes with diverged co-expression that could explain these differences, we first identified orthologs with conserved co-expression within, but not between, the two lineages (Fig. 4A). This analysis resulted in 1,811 pairs of dicot-/conifer-specific cliques from 294 orthogroups (Tab. S4). These included genes enriched for pathways related to carbohydrate metabolism, nitrogen signaling, and intracellular trafficking (Tab. S5), suggesting divergent strategies in resource allocation, metabolic coordination, and cellular remodeling during secondary growth. One illustrative example is the ortholog of *phragmoplast orienting kinesin 2* (*POK2*) (Fig. 4B). Phragmoplasts are specialized cytoskeletal structures that form during the late stages of plant cell division to build the cell plate - a partition between two daughter cells that precedes primary cell wall (PCW) deposition. While *POK2* showed peak expression in the cell division zone in both lineages, consistent with its likely role in maintaining proper cell plate formation and alignment, it also displayed a second expression peak during secondary cell wall (SCW) formation in conifers only (Fig. 4B). It is tempting to speculate that this unexpected pattern may reflect a conifer-specific role for cytoskeletal elements in guiding localized wall deposition processes in tracheids, such as pit membrane thickening or remodeling.

**Fig. 4.**
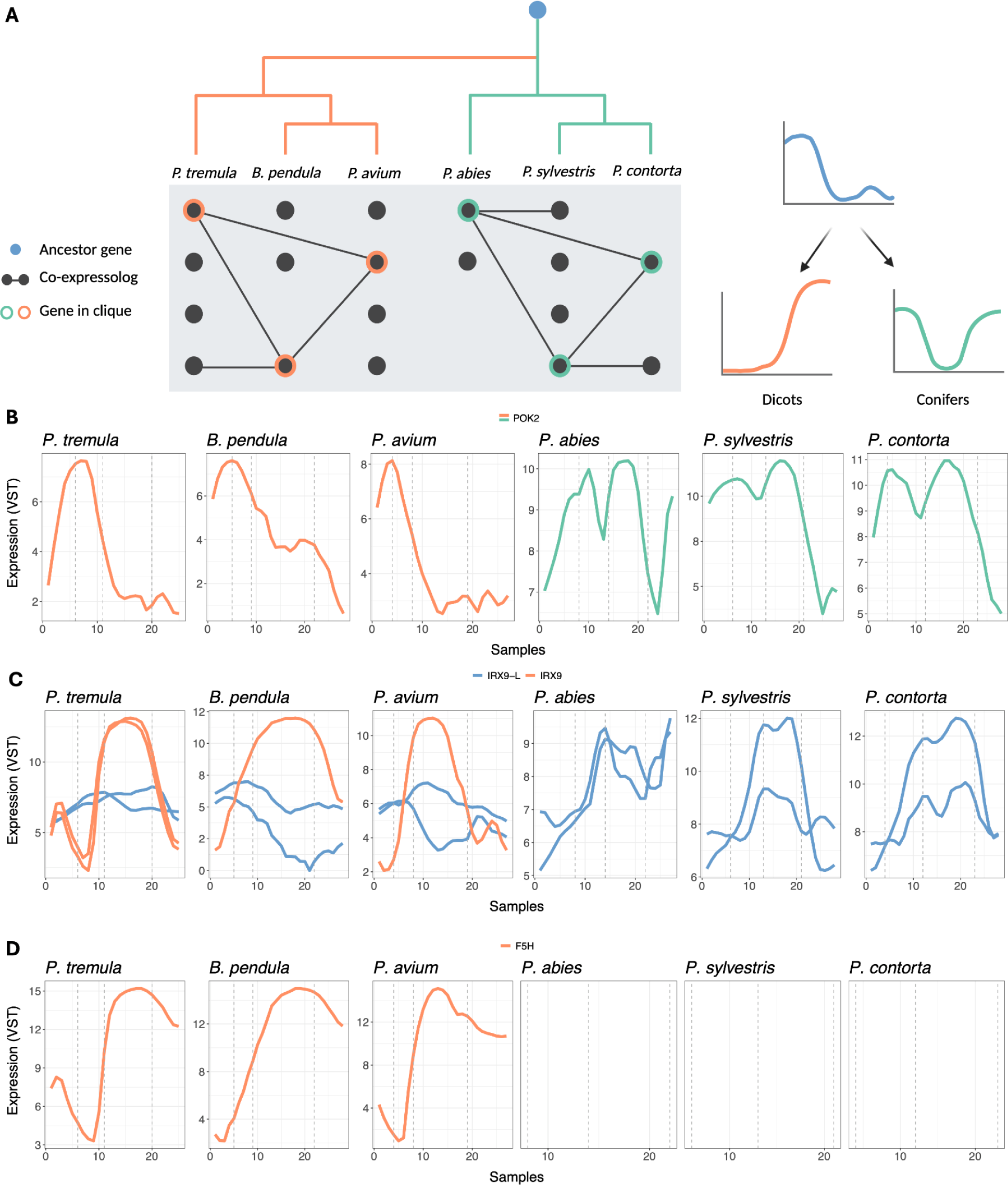
Genes with diverged expression between dicots and conifers. (A) Conceptual figure of how we selected cliques of genes with conserved co-expression (i.e. co-expressologs) within dicots and within conifers, but not between dicots and conifers. (B) Gene with diverged expression between dicots and conifers: phragmoplast orienting kinesin 2 (POK2). The gene forms a dicot- and a conifer-specific clique, but does not have conserved expression across dicots and conifers. (C) Orthologs of IRREGULAR XYLEM 9 (IRX9) form two overlapping dicot-specific cliques (same gene in B. pendula and P. avium). The paralog in Arabidopsis (IRX9-L) has two orthologs in each of the six tree species. (D) Orthologs of FERULIC ACID 5-HYDROXYLASE (F5H) form a unique dicot-specific clique.

We also analyzed the 2,093/3,640 orthogroups (Fig. 2A) that were either lineage-specific or contained genes only expressed in wood within one lineage, identifying 1,160 dicot-specific and 3,045 conifer-specific unique cliques. Dicot-specific cliques included several genes relevant to wood formation such as *xylem serine peptidase 1* (*XSP1*), *IRREGULAR XYLEM 8* (*IRX8*) and *IRX9*, *VND-interacting 2* (*VNI2*) and *FERULIC ACID 5-HYDROXYLASE* (*F5H*). *IRX8* and *IRX9* are key xylan end sequence- and backbone-biosynthetic genes, respectively. *IRX9* and its paralog *IRX9L* have been proposed to form secondary and primary cell wall xylan synthase complexes, respectively^29^. Our data support that *IRX9* genes have a specialized role in SCW biosynthesis in dicots, with a clear peak during SCW formation, while *IRX9L* genes exhibit a more general expression pattern during xylogenesis similar to that of pectin and xyloglucan biosynthetic genes (Fig. 4C). Syringyl lignin (S-lignin) is a hallmark of dicot wood, contributing to a more flexible and less condensed lignin polymer compared to the guaiacyl (G) lignin-dominated structure in conifer wood, with *ferulate 5-hydroxylase* (*F5H*) catalyzing the rate-limiting step in S-lignin biosynthesis^30^. The dicot-specific expression of *F5H*, with a profile resembling that of SCW cellulose synthase genes (Fig. 4D), is congruent with the absence of S-lignin in conifers. Conifer-specific cliques, on the other hand, included genes involved in pathogen defense (TMV resistance protein) and pentatricopeptide repeat (PPR)-containing proteins, which are responsible for RNA editing and that have experienced multiple lineage-specific duplications in conifers^31^.

### Acetylation of cell wall polysaccharides across dicots and conifers

Acetylation plays a crucial role in regulating cell wall properties by modifying the hydration status, enzymatic accessibility, and intermolecular interactions of cell wall polymers, which influence wall porosity, mechanical strength, and resistance to degradation. Trichome Birefringence-Like (TBL) proteins include key O-acetyltransferases responsible for acetylating specific cell wall polysaccharides during their biosynthesis in the Golgi, as well as recently identified acetyl esterases that deacetylate specific polysaccharides in cell walls^32^. Given the difference in hemicellulose acetylation and composition between angiosperm and conifer wood, differences in TBL family structure and activity are expected. In angiosperms, glucuronoxylan is the dominant hemicellulose, whereas (galacto)glucomannan is less abundant and both of these hemicelluloses are acetylated^33^. In contrast, acetylated (galacto)glucomannan is the main hemicellulose in conifers whereas arabinoglucuronoxylan is less abundant and non-acetylated^34^. Both lineages also contain low amounts of xyloglucan, which is known to be acetylated in angiosperms^35^, whereas its acetylation in conifers has not, to our knowledge, been analyzed.

Analyzing the phylogeny of different orthogroups of the *TBL* family (Fig. S4), we discovered that *TBLs* responsible for xylan (*TBL 3* + *28-35*) and xyloglucan (*TBL 19-22* + *27*) acetylation lack orthologs in conifers. We identified nine unique *TBL*-cliques containing genes with conserved co-expression across all six tree species (conserved) and five unique cliques specific to dicots (dicot-specific). Among the conserved *TBLs* (Fig. 5A), *TBL12* and *TBL44/PMR5*, characterized as both rhamnogalacturonan I (RGI)- and homogalacturonan (HG)-specific^36^ and only HG-specific^37^, respectively, showed expression peaks in the expansion zone, consistent with their role in pectin modification during primary cell wall formation (PCW). *TBL25,* which is specific to glucomannan^38^, peaked during SCW deposition and declined as lignification began, consistent with its role in glucomannan acetylation. Interestingly, *TBL37*, which is a jasmonate-induced *TBL* member responsible for non-specific hemicellulose acetylation^39^, showed a distinct pattern between the lineages: in dicots, it peaked during the SCW zone, whereas in conifers, its expression peaked later in the maturation zone, potentially indicating differences in jasmonate signaling between dicots and conifers.

**Fig. 5.**
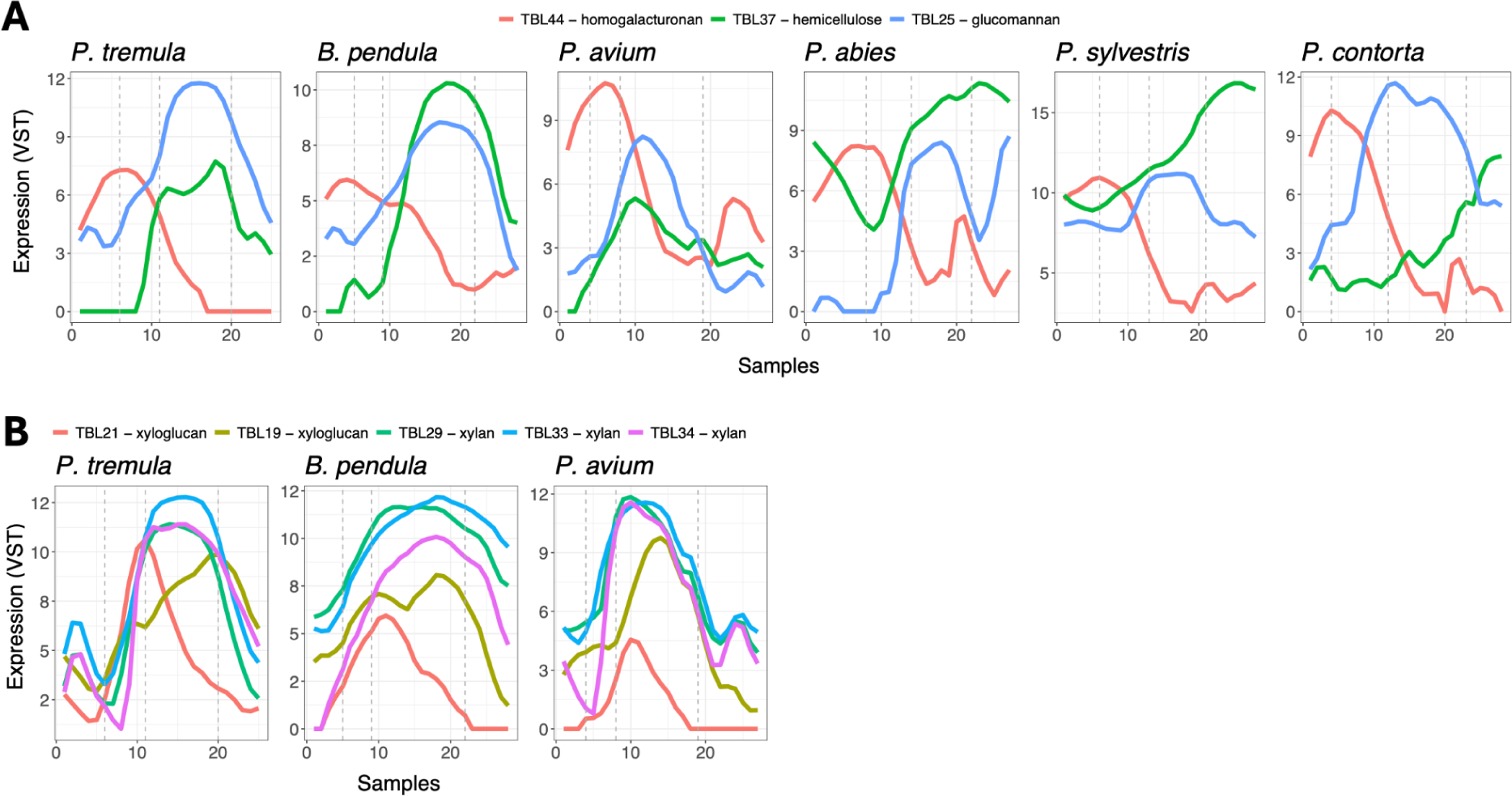
Trichome Birefringence-Like (TBL) gene expression. (A) TBLs with conserved expression across dicots and conifers. (B) Dicot-specific TBLs.

Dicot-specific *TBL*s included those targeting the xyloglucan backbone (*TBL19*, *TBL21*^40^) and xylan (*TBL29*, *TBL33*, *TBL34*^41–45)^ (Fig. 5B). As expected, *TBL21* showed increasing expression during cell expansion, consistent with a role in xyloglucan in PCW. In contrast, *TBL19* peaked much later, near the end of SCW formation, suggesting a possible role in xyloglucan biosynthesis in the protective layer that is formed at later stages of cell wall differentiation after SCW deposition^34^. The xylan-associated *TBL*s showed clear expression peaks during SCW deposition, supporting their role in xylan acetylation.

Taken together, our findings suggest that core TBL functions in pectin and glucomannan acetylation are conserved across dicots and conifers, whereas xylan and xyloglucan acetylation is specific to dicots. The absence of xyloglucan-related TBLs and the presence of pectin-related TBLs in conifers highlight the need for further analyses of xyloglucan and pectin acetylation in conifers.

### The regulatory network underlying wood formation

Our spatially resolved transcriptomes represent an excellent resource for gaining further insight into the regulatory logic controlling wood formation that is hard-wired in the genomes of trees. In Arabidopsis, the regulatory network controlling SCW formation is organized into three hierarchical layers of transcription factors with overlapping functions: a top layer of NAC master regulators *VASCULAR-RELATED NAC-DOMAIN* (*VND1–7*) and *NAC SECONDARY WALL THICKENING PROMOTING FACTORS* (*NST1–3*); a middle layer consisting of R2R3-MYB transcription factors MYB46 and MYB83; and a bottom layer that includes additional MYBs and *SECONDARY WALL-ASSOCIATED NAC DOMAIN* (*SND2 and 3*)^46,47^.

In Arabidopsis, VNDs control the formation of the SCW in vessels while NSTs control SCW formation in fibers^46^. We found that orthologs of VNDs exhibited conserved expression patterns across all six tree species, with a distinct expression peak during SCW formation (Fig. 6A). In dicots, orthologs of NSTs displayed a similar expression peak during SCW formation, while in conifers the single NST-copy had a pronounced dip in expression at this stage (Fig. 6B). Consistent with what was previously observed in a pairwise comparison of *P. tremula* and *P. abies*^12^, our data thus provide strong evidence that NSTs have diverged in expression between dicots and conifers.

**Fig. 6.**
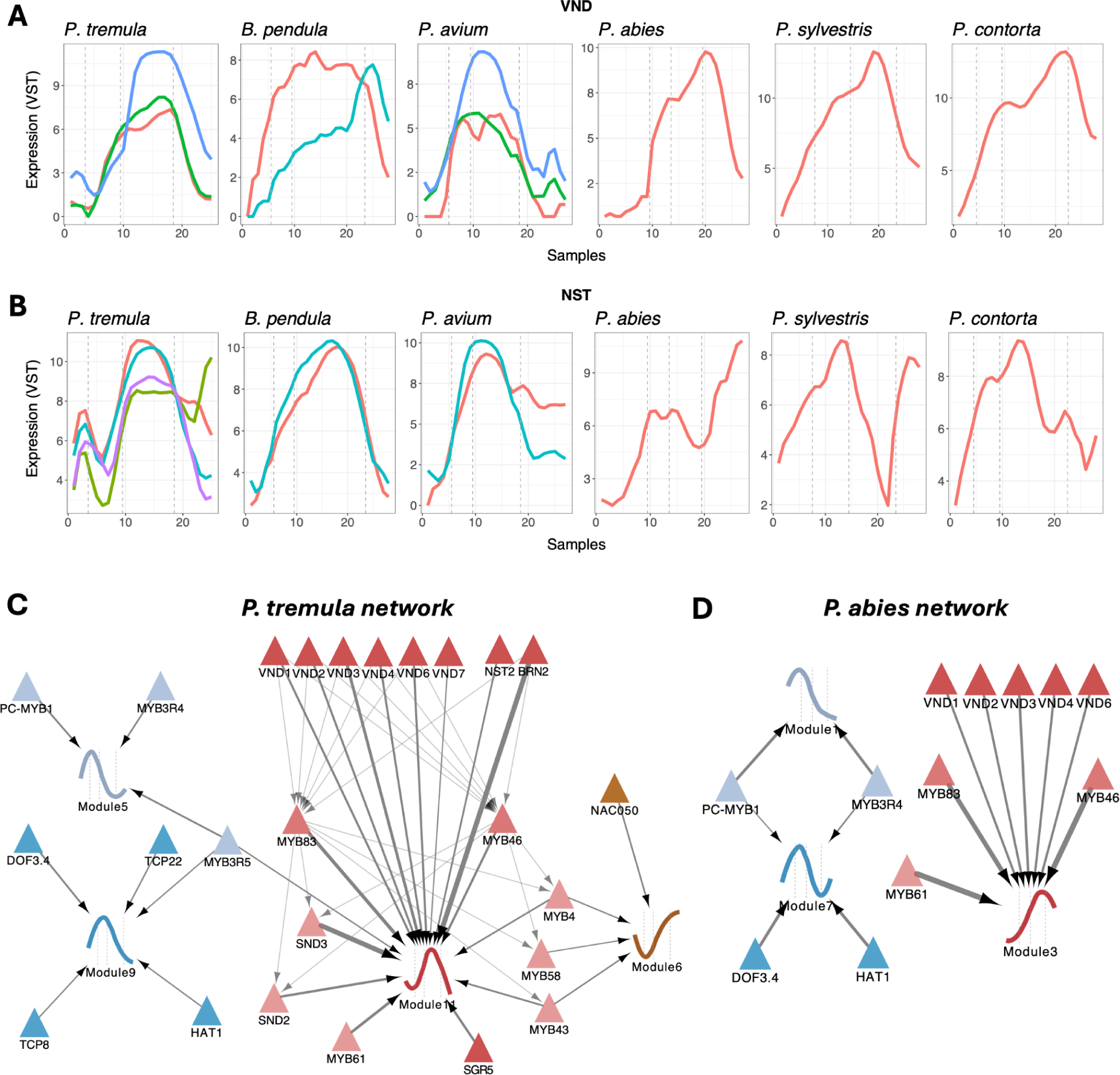
The Gene Regulatory Network (GRN) of Populus tremula and Picea abies. (A) Conserved cliques of VND master regulators. (B) Expression profiles of NST master regulators. (C,D) The P. tremula/P. abies regulatory network restricted to the modules of the five marker genes (displayed using the expression profile of the central gene in the module) and a selected set of top scoring characterized regulators (triangles). Neither network contained regulators of the phloem module, while the secondary cell wall (SCW) and cell death modules were merged in P. abies. The width of the links indicates the strength of the inferred regulatory interaction, combining the strength of the enrichment of the TF motif in the module and the strength of the expression similarity between the TF and the module. Regulators are colored according to the module with the strongest inferred regulatory interaction. SCW regulators are organized into a hierarchy of three layers (indicated by transparency), and links between layer 1 and 2, and layer 2 and 3 are based on the occurrence of individual motifs. The orthogroup containing Arabidopsis NST 1-3 also included BRN 1-2 (BEARSKIN 1-2) and SMB (SOMBRERO) (Fig. S5). While the closest orthologs of BRNs and SMBs in our six species were not expressed in wood, the BRN2- and NST2-binding motifs were highly similar and, in our network analysis, linked to the same NST gene in P. tremula (Potra2n2c4785, purple gene in panel B) and P. abies (PA_chr11_G001721, single gene in panel B). We therefore included BRN2 in the network as a potential binding site for NST.

To gain further insight into the regulatory role of these SCW regulators, we generated Assay for Transposase-Accessible Chromatin using Sequencing (ATAC-seq) data from developing wood (xylem scrap) in *P. tremula* and *P. abies*. We then mapped characterized DNA sequence patterns recognized and bound by plant TFs (i.e. TF DNA binding motifs) to open chromatin regions in the two tree genomes. These putative binding sites were linked to the closest gene (target) and the motif was associated with a co-expression network module (i.e. a set of co-expressed genes) if its target genes were statistically significantly enriched in the module. Finally we used orthogroups and co-expression to assign a specific TF gene to the motif (see Methods). The resulting robust TF-module networks linked 175/165 TFs to 13/8 modules containing 8,228/5,489 genes in *P. tremula* and *P. abies*, respectively (Tab. S6). As expected, the regulatory network did not contain phloem-specific regulation as phloem cells were not well-represented in the ATAC-seq samples.

In *P. tremula*, the co-expression module containing the SCW *CesA* genes (module 11) was strongly preferentially associated with TFs known from the Arabidopsis SCW regulatory network, including the top layer master regulators VND and NST, the middle layer MYB46 and MYB83 and several bottom layer MYBs as well as SND2 and SND3 (Fig. 6C). Three bottom layer MYBs were also predicted to regulate the mature xylem module containing the cell death marker gene (module 6), indicating a previously unknown mechanism coordinating SCW formation and cell death/lignification (Fig. 6C). Several of the core SCW master regulators including VNDs, MYB46 and MYB83 were also associated with the corresponding SCW module in *P. abies* (module 3, Fig. 6D), thus indicating strong purifying selection on their binding sites and a common origin of the regulatory network of wood formation.

Next, we specifically investigated whether the top regulators of the SCW *cesA* genes differ between dicots and conifers. In *P. tremula*, *SND3*, *VND3*, and *NST2* emerged as the strongest candidates, showing both high expression correlation with the SCW *cesAs* (R > 0.85) and predicted binding sites associated with all five SCW *cesAs* (Tab. S6). In *P. abies*, *MYB83* was the strongest candidate, with high correlation (R = 0.88) and predicted binding sites associated with all three SCW *cesA* genes, followed by *MYB46* and several *VNDs* (Tab. S6).

Taken together, the structure of the *P. tremula* regulatory network mirrors that of Arabidopsis, adopting a layered architecture with feed-forward loops linking VNDs/NSTs, MYBs/SNDs, and biosynthetic genes. In contrast, the *P. abies* network is notably flatter: VNDs and MYBs act more in parallel, with MYBs showing the strongest predicted regulatory interactions with biosynthetic targets, while NSTs appear unlikely to serve as primary regulators of SCW *cesAs* in this lineage. These findings provide new insight into the regulation of SCW formation in conifers and highlight specific *P. abies* MYBs as promising targets for biotechnology and breeding efforts to tailor wood properties.

### Resource for wood biology

Our datasets provide an unprecedented resource for wood biology, enabling the scientific community to address long-standing questions and generate insights beyond the scope of this paper. We have made all data available at PlantGenIE.org.

## Materials and methods

### Sampling

Wood samples from *Betula pendula*, *Pinus sylvestris*, and *Picea abies* were obtained at a clonal field trial in Skogforsk, Sävar (63.895648°N, 20.549005°E). *Betula pendula* samples from genotype 18 were taken on July 8th, 2020, between 10:30–13:30. *Pinus sylvestris* samples from genotype 29 were taken on July 8th, 2020, between 13:30–14:30. *Picea abies* samples from genotype K27-2125 were taken on July 10th, between 10:30–12:00. Wood samples from *Pinus contorta* from genotype 86 were obtained at the clonal field trial in Fagerlund (plot 7114, 64.083393°N, 19.793286°E), on July 9th, 2020, between 10:30–12:00. Wood samples from *Prunus avium* from genotype S21-K946 6006 were obtained from Skogforsk, Ekebo (55.950860°N, 13.112653°E), on June 28th, 2021, between 10:00–12:00. No obvious abiotic or biotic stresses were observed at the time of sampling.

Rectangular wood blocks were collected from each tree, immediately placed in dry ice boxes and stored at -70 °C. Prism-shaped pieces, containing the main layers of wood from cortex to mature xylem, were cut from the wood blocks using an electric saw. These pieces were placed into a cryo-microtome and multiple longitudinal sections 15 µm thick were cut from each layer ^48^. Only pieces with clearly discernible developmental layers were used. During the cutting process, cross sections were frequently prepared and studied under a light microscope to characterize the different tissue each section originated from. Between 120 and 150 longitudinal sections were prepared from each prism-shaped piece, covering the entire current year’s growth and the four developmental layers: phloem, cambium, early/developing xylem and mature xylem. Each section was stored in a separate 1.5 ml Eppendorf tube and stored at -80 °C. This was repeated three times for a total of three replicates per tree.

### RNA extraction

For each replicate tree, sections were pooled together as shown in Tab S1 and thawed in QIAzol lysis reagent (Qiagen). Total RNA was extracted separately from each section pool using a protocol adapted from Chang *et al.* (1993)^49^ and the Qiagen RNeasy Kit (Qiagen, Hilden, Germany), with modifications following Schrader *et al.* (2004)^50^. Extraction buffer containing 2% CTAB, 100mM Tris-HCl (pH 8.0), 25 mM EDTA, 2.0 M NaCl, and 0.5g/L spermidine was prepared and pre-warmed. The Eppendorf tubes with section tissue samples were taken from -80 °C storage onto dry ice. At the start of the extraction protocol, the extraction buffer was added into the Eppendorf tubes and the tissue samples were dissolved. Chloroform:isoamylalcohol was added and phases were separated by centrifugation at 10 000 rpm for 10 minutes. The supernatant was transferred to a new tube without disturbing the interphase, and the extraction step was repeated. RNA precipitation was carried out by adding 2.0 volume ethanol and incubating for 1 hour at 4°C, followed by centrifugation at 14 000 rpm for 20 minutes at 4°C, resulting in a pellet.

The extraction continued with the Qiagen RNeasy Mini Kit. The RNA pellet was dissolved in 350 µL of RLT buffer from the Qiagen kit, added 1 volume of 70% ethanol, transferred to the column tube and spun at 10 000 rpm for 15 seconds. The column was then washed with 350 µL of buffer RWT and spun the same way.

A DNase treatment was performed by mixing 10 µL of DNase I stock solution with 70 µL of buffer RDD, which was loaded onto the column and incubated at room temperature for 30 minutes. The column was washed and spun once with 350 µL of buffer RWT and twice with 500 µL of buffer RPE. The column was dried by centrifugation at 12 000 rpm for 5 minutes. The final RNA was eluted adding 35 µL of RNase-free water and spinning at 10 000 rpm for 1 minute. Total RNA from each section pool was quantified using the RNA High sensitivity programme in the Qubit 2.0 fluorometer (Life Technologies, Carlsbad, California, USA).

### RNA library preparation and sequencing

The Universal Plus mRNA-Seq with NuQuant protocol (Tecan, Männedorf, Switzerland) was used to prepare cDNA libraries from the extracted RNA of each of the section pools according to the provided protocol. Following adapter ligation, the cDNA libraries were amplified by PCR and purified using Agencourt AMPure XP beads (Beckman Coulter, Brea, California, USA).

Final concentration was measured using Qubit RNA High sensitivity and quality was studied with the Agilent Bioanalyzer 2100 (Agilent Technologies, Santa Clara, California, USA) using Pico chips.

The libraries were sequenced on NovaSeq PE150 to a depth of 20 million pair-end reads. Raw RNA-Seq reads were submitted to the European Nucleotide Archive (http://www.ebi.ac.uk/ena/) under accession numbers ERP016242 (*Populus tremula*), E-MTAB-14145 (*Betula pendula*), E-MTAB-14172 (*Prunus avium*), E-MTAB-14173 (*Picea abies*), E-MTAB-14175 (*Pinus sylvestris*) and E-MTAB-14176 (*Pinus contorta*).

### Read mapping and expression quantification

The resulting reads were quality checked and filtered using a publicly available preprocessing pipeline for RNA-seq data implemented at the Umeå Plant Science Centre (v0.4.0, https://github.com/UPSCb). First, individual quality reports were generated for each sample using FastQC v0.11.9^51^ and summarized for each species with MultiQC v1.11^52^. We filtered out ribosomal RNA with SortMeRNA v4.3.4^53^, and used Trimmomatic v0.39^54^ to remove adaptors and trim reads with a 5 bp sliding window and a quality threshold of 30. For each species, filtered and trimmed reads were mapped to the reference with the alignment-free quantification tool Salmon v1.6.0^55^. We enabled options for modeling position-specific fragment start distribution, sequence-specific bias correction, and GC-content bias correction. For the dicot species, we additionally used a decoy-aware index to improve mapping accuracy; this was not feasible for the conifer species due to the large genome sizes. The following CDS and genomes versions were used in the analyses: *Populus tremula* v2.2^56^, *Betula pendula* v1.2 (earlier genome version used after recommendation from the authors^57^), *Prunus avium* v2.0^58^, *Picea abies* v2.0, and *Pinus sylvestris* v1.0 (used for both *P. sylvestris* and *P. contorta*)^26^.

For each tree species, read counts were subjected to a moving average smoothing (within each replicate tree) before normalization using the Variance Stabilizing Transformation (VST) as implemented in DESeq2^59^. Samples were inspected using hierarchical clustering and principal component analysis (PCA), and outliers were removed iteratively (i.e. normalization was redone, removed samples are marked in Tab. S1: 18 samples in *B. pendula* and 15 samples in *P. abies*). Genes with VST normalized expression values > 3 in two samples in three replicate trees (two replicate trees in *B. pendula* and *P. abies* due to the large number of samples removed from one tree) were defined as expressed. Hierarchical clustering was conducted using the R-function *hclust* with parameter *method="ward.D2"* and Person correlation. PCA was done using the R-function *prcomp*.

### Orthogroups

Orthogroups were inferred using OrthoFinder v2.5.2^60^, based on the longest protein-coding sequence per gene from annotated genomes of 27 plant species, and incorporating a rooted species tree obtained from TimeTree. We used both standard orthogroups (OGs) and high resolution Phylogenetic Hierarchical Orthogroups (HOGs: N1 with *Physcomitrella patens* as the outgroup). Statistics reported in the paper are for HOGs. The gene content of the orthogroups were visualized using the *UpSetR* package^61^.

### Gene co-expression analysis

#### Identifying co-expressologs using ComPlEx

Co-expression networks were generated for each species using Pearson’s correlation and Mutual Rank (MR) normalization, as implemented in the popular PlaNet method^62^. The networks were constrained to include only the top 3% most strongly co-expression links. For each of the 15 pairs of species, co-expression neighborhoods of orthologs were compared using the ComPlEx method^23^. Briefly, given an ortholog pair A-B, the co-expression neighbors of ortholog A were mapped to their ortholog(s) in the network of ortholog B, and the overlap between the mapped neighbors of A and the neighbors of B was assessed using the hypergeometric test. Similarly, the neighbors of B were mapped to the network of A resulting in two p-values, one using the network of ortholog A as the background and one using the network of ortholog B. We required both False Discovery Rate-corrected p-values to be below 0.1 to classify the pair as a *co-expressolog*.

#### Sample comparison

The expression similarity of samples coming from different species were visualized using sample correlation heatmaps. For each species pair, the expression similarity of each sample-pair was quantified by computing Pearson’s correlation of the expression values of all co-expressologs. Since a gene could occur in multiple co-expressolog-pairs, duplicates were removed, retaining the co-expressolog pair with the lowest FDR-value. Heatmaps were generated using the ComplexHeatmap package.

#### Identifying cliques

For each orthogroup, a network was created where orthologs were connected by links with weights set to the largest co-expressolog FDR-value. Cliques were then identified using the Igraph package and the function *max_cliques*. First, genes conserved across all six species were identified as six-member cliques (i.e. including one gene from each of the six species). We included both “complete conserved cliques” requiring all 15 ortholog-pairs to be co-expressologs (FDR < 0.1) and “partial conserved cliques” requiring at least 11 ortholog-pairs to be co-expressologs (with all gene pairs required to have FDR < 0.9). To account for missing gene annotations in individual species, we also classified five-member cliques as “partial conserved cliques” if all ortholog-pairs were co-expressologs. Differentiated cliques were identified as six-membered cliques where all three lineage-specific ortholog-pairs were co-expressologs and no more than four inter-lineage pairs were co-expressologs. Lineage-specific cliques were identified from orthogroups containing only dicot or conifer expressed genes, and were required to be complete, three-membered cliques.

Conserved cliques (complete and partially significant) were visualised using the ComplexHeatmaps package using correlation as a distance metric and clustering using the ward.D2 method. Scaling and centering of gene expression was performed individually for each species prior to clustering.

#### Enrichment analysis

Gene ontology enrichment analysis was performed for the conserved and differentiated gene sets as well as for clusters of conserved genes using the GSEABase and GOstas packages. The analysis was based on annotated *Populus tremula* genes downloaded from plantgenie.org. A Fisher exact test was performed using the *GSEAGOHyperGParams* function setting ontology to biological process (BP) and using a p-value cutoff of 0.05. The significant GO terms were summarised using Revigo^63^ and visualised as treemaps. Segment sizes were based on the *values* column containing -log10-transformed enrichment p-values.

### Regulatory networks analysis

#### Transcriptional module detection

To infer robust regulatory networks, we based our transcription factor (TF) motif analysis on modules of co-expressed genes rather than individual genes. Modules were detected using a coverage algorithm: We iteratively selected the most highly connected genes in the co-expression network (centroids) with neighbors (top 1%) not overlapping previously selected neighborhoods. Genes were assigned to their closest centroid if the link was in the top 3%.

#### Open chromatin regions: Nuclei for ATAC-sequencing

The scrapings of developing xylem from aspen and spruce trees were powdered by grinding in liquid nitrogen. The nuclei were purified using a Percoll gradient method^64^. In brief, 2g each of grounded tissues in were resuspended in 20 ml of 1X nuclei isolation buffer (NIB) [10 mM MES-KOH (pH 5.4), 10 mM NaCl, 10 mM KCl, 2.5 mM EDTA, 250 mM sucrose, 0.1 mM spermine, 0.5 mM spermidine, 1 mM DTT] supplemented with 2 % (w/v) Polyvinylpyrrolidone-40 (PVP-40, Sigma-Aldrich) and 0.1 % Protease inhibitor cocktail (Roche) just before extraction. The homogenates were gradually filtered through two layers of pre-wetted cheesecloth and one layer Miracloth. In filtrate, Triton-X100 was added in a dropwise manner to make the concentration 1% (v/v) and incubated for 20 min at 4°C under gentle rotation. The homogenates were centrifuged 2000 x g for 10 min at 4°C and pellets were resuspended in 5 ml 1X NIB with help of paint brush. The crude preparation of nuclei suspension was carefully loaded on the top of the pre-assembled density gradient (3 ml 60% Percoll in 1x NIB and 3 ml of 2.5 M Sucrose) by slowly releasing the solution onto the side of the tube above the 60% Percoll layer with the help of Pasteur pipette. The assembled gradient is subject to centrifugation in a swinging bucket rotor at 1200 × g for 30 min at 4°C without breaks. The rest of the nuclei purification steps were kept as described for *Rosaceae* in^64^. The DAPI (0.5 μg/ml) stained nuclei were counted using Neubauer chamber (Sigma-Aldrich) under Axioplan 2 (Ziess) microscope with DAPI filter, and for Aspen 40000 and for Spruce 30000 nuclei were used for ATAC -seq library preparation.

The ATAC -seq libraries were prepared after optimising the amount of Tn5, number of nuclei and time of incubation for each species as explained in Picelli *et al.* (2014)^65^ for in-house produced Tn5. Further 40000 and 30000 nuclei from aspen and spruce were tagmented for 30 min at 37oC with 3µl of adapter loaded-Tn5, tubes were flipped after each 10 min of incubation. The tagmented DNA was purified using MinElute PCR Purification Kit (Qiagen Cat. No. / ID: 28004) and eluted in 12µl nuclease free water. The eluted DNA was used for sequencing library preparation using indexing primers Illumina Nextera Library Prep Kits and the size selected libraries were sequenced on NovaSeq 6000 in pair end mode.

#### Aspen and spruce ATAC-seq pre-processing

Trimmomatic v0.39^54^ was used to perform quality trimming on raw paired-end reads with the setting: ‘ILLUMINACLIP:$TRIMMOMATIC_HOME/adapters/NexteraPE-PE.fa:2:30:10:1:TRUE SLIDINGWINDOW:5:20 MINLEN:38’. Trimmed reads were aligned to the *P. abies* and *P. tremula* reference genomes for their respective datasets using Bowtie2 v2.4.5^66^ with the setting ‘--very-sensitive --dovetail --maxins 1000’. Alignments were filtered using Samtools v1.16^67^ with ‘-f 3 -F 12 -q 20’ to retain high-quality, properly paired reads (MAPQ ≥ 20), followed by PCR duplicate removal with Samtools markdup. Peaks were called using MACS3 v3.0.0b1^68^ with the setting ‘--format BAMPE’ and genome sizes of ‘--gsize 17.7e9’ for *P. abies* and ‘--gsize 3.8e8’ for *P. tremula*. The ‘--keep-dup all’ option was used, as PCR duplicates had already been removed prior to peak calling.

#### Inference of Gene Regulatory Network (GRN)

We compiled 808 TF motifs from the JASPER^69^ and PlantTFDB databases^70^, and linked an Arabidopsis TF to each motif using a combination of BLASTp and the name of the motif. The motifs were mapped to open chromatin regions in *P. tremula* and *P. abies* using the Find Individual Motif Occurrences (FIMO) tool in the MEME suite, version 5.5.5^71^. Motifs were assigned to the closest gene within 10 kbps (target genes), and a motif was assigned to a network module if the target genes of that motif were enriched in the module (Fisher exact test, FDR-corrected p-value < 0.05). Given that a motif was enriched in a module, we searched the orthogroup of the associated Arabidopsis TF for the *P. tremula*/*P. abies* TF with the highest expression correlation to the average expression profile of the module. The *P. tremula*/*P. abies* TF were assigned to the motif/module if the Pearson correlation > 0.7 and the FDR-corrected p-value < 0.05. The TF-motif-module associations were scored as *-log10(Motif enrichment p-value) * Expression correlation*. TF-TF linked was inferred based on individual motif matches. GRNs were visualized by Cytoscape, v3.10.3^72^.

## Code availability

All code generating results and figures in this article can be found at: https://github.com/ellendim/EVOTREE.

## Acknowledgements

We thank the UPSC bioinformatics facility for support. We thank Skogforsk for locating suitable material and enabling sampling of all species. We thank Mikko Luomaranta and Kathryn Robinson for assistance with sampling.

## Author contributions

ER and ZCL prepared the cryo-sections. ER, SB, VK and ZCL prepared the RNA-seq libraries. SB did RNA-seq read mapping. EDC performed the RNA-seq data analysis. SF performed the gene regulatory network analysis. VK prepared the ATAC-seq libraries. TAK performed the ATAC-seq data analysis. JH generated the orthogroups. HT and EJM contributed significantly to data interpretation. NRS and TRH conceived the study and obtained funding. All authors read and approved the final manuscript.

## Supplemental figures

Figure S1. Heatmaps for each species. For each species, a heatmap of all expressed genes (rows) and all retained samples (column) are shown. The expression values of each gene are scaled and colored from blue (lowest expression) to red (highest expression). Each replicate tree is separated by a black line. The horizontal color bar marks four sample clusters largely coinciding with the developing phloem (green), the cell expansion zone (blue), the secondary cell wall zone (red) and the maturing xylem zone (brown). The vertical color indicates gene clusters.

Figure S2. Comparisons of six wood transcriptomes. (A) The number of orthogroups with at least one co-expressologs - for each of the 15 species pairs. The total number of co-expressologs are in parentheses. (B) The distribution of correlations between samples from different species pairs: three dicot pairs, three conifer pairs and nine inter-lineage pairs (dicot-conifer pairs).

Figure S3. H3.1 and H3.3 phylogeny. The phylogeny of the orthogroup (OG0000095) contained all Arabidopsis H3.1 (HTR1, 2, 3, 9 and 13) and H3.3 (HTR4, 5 and 8) genes.

Figure S4. Trichome Birefringence-Like (TBL) phylogeny. The common phylogeny of all genes in orthogroups containing Arabidopsis TBL genes.

Figure S5. NAC phylogeny. The phylogeny of NAC master regulators.

## Supplemental tables

**Table S1. Cryosections and pooled samples for each species**. For each species and each replicate tree, a diagram is showing the original cryosection (Section), the sections pooled into samples for RNA-seq (Sample) and the annotated wood tissue (Tissue: Phloem, Cambium, Xylem). Samples excluded from downstream analysis are marked in the Outlier-column.

**Table S2. Gene expression data sets**. For each species, a table is provided with genes as rows, samples as columns and VST-normalized expression as values.

**Table S3. Co-expressologs**. For each species pair, ortholog pairs with significantly overlapping co-expression neighborhoods (i.e. co-expressologs) are listed.

**Table S4. Cliques.** Tables of different types of cliques conserved across all species (Conserved), conserved within dicots and within conifers, but not across the two lineages (Differentiated) and conserved specifically in dicots or conifers (Specific). Each table exists in two versions: all cliques and unique cliques. Cliques computed for (A) high resolution Phylogenetic Hierarchical Orthogroups (HOGs) and (B) standard orthogroups (OGs).

**Table S5. Gene function enrichment**. Enriched Gene Ontology (GO) Biological Processes among genes in the different clique types in Tab. S4A.

**Table S6. Gene regulatory network**. Tables containing the inferred regulatory network for *P. tremula* and *P. abies*, as well as the Position Frequency Matrices (PFMs) for the analyzed transcription factor motifs.

## Supplementary material

Supplemental figures and tables as well as high-resolution figures can be downloaded at: https://tinyurl.com/SupplementWoodRegulomics.

